# VIPR: Vectorial Implementation of Phase Retrieval for fast and accurate microscopic pixel-wise pupil estimation

**DOI:** 10.1101/2020.01.02.892661

**Authors:** Boris Ferdman, Elias Nehme, Lucien E. Weiss, Reut Orange, Onit Alalouf, Yoav Shechtman

## Abstract

In microscopy, proper modeling of the image formation has a substantial effect on the precision and accuracy in localization experiments and facilitates the correction of aberrations in adaptive optics experiments. The observed images are subject to polarization effects, refractive index variations and system specific constraints. Previously reported techniques have addressed these challenges by using complicated calibration samples, computationally heavy numerical algorithms, and various mathematical simplifications. In this work, we present a phase retrieval approach based on an analytical derivation of the vectorial diffraction model. Our method produces an accurate estimate of the system phase information (without any prior knowledge) in under a minute.

## 1. Introduction

Attaining phase information in diffraction experiments is of great practical interest; however, intensity measurements, i.e. images, do not explicitly capture complex values. This hidden information can be estimated by Phase Retrieval (PR) [1] – namely, recovery of a complex signal from intensity measurements of its Fourier transform, which has therefore been employed in a wide variety of applications, e.g. X-Ray crystallography [2], astronomy [3], optical ptychography [4,5], adaptive optics [6,7], and the subject of this work: localization microscopy [8,9].

In a typical 2D localization microscopy experiment, the localization step is performed by fitting images of well-separated emitters with a known Point-Spread Function (PSF), e.g. an Airy disc. The reliance on precise knowledge of the PSF is highlighted in single-molecule localization microscopy (SMLM), where the Signal-to-Noise Ratio (SNR) is limited. Acquiring additional information can be achieved by PSF engineering, where the PSF shape is intentionally perturbed to encode the desired information. This approach has been used for 3D measurements, e.g. astigmatism [10], Tetrapod [11] and Double Helix (DH) [12] PSFs; encoding emitter wavelength [13–15] and molecular orientation [16]. A closely related application is adaptive optics, which employs additional optical elements to counteract aberrations present in the imaging system that distort the PSF. In all mentioned cases, PR can be used to correct the theoretically simulated PSFs or to design the phase mask which creates them [17].

In PR, the complex signal is numerically estimated from the magnitude of its Fourier transform. Any employed algorithm requires the nonobvious selection of a penalty function, and must overcome degeneracies in the solution, unstable derivatives, non-convex optimization, and more [18]. So far, PR methods have made use of the Iterative Gerchberg-Saxton(GS) Algorithm [19] and its variations [20], estimation over the Zernike basis of aberrations [21–23], and pixel-wise numerical gradient calculations [24]. The first consideration of these algorithms is the tradeoff between computational efficiency and model accuracy; namely, a limited number of parameters must be chosen to describe the phase mask and various model approximations are employed. To reduce computational burden, the mentioned discrete approaches employ coarse graining in the physical space, while the methods that use a polynomial basis, e.g. Zernike polynomials, reduce the number of parameters by modeling only the most common optical aberrations.

A second consideration is the selection of an optical model. Thus far, most methods employ the scalar diffraction model [21,25]. The scalar model is more computationally efficient to simulate than the vectorial model; however, this approximation deteriorates for emitters near the coverslip, precisely where most calibration measurements are performed. While in some cases, PR can be performed within a biological sample (away from the coverslip), which would make the scalar model appropriate [26], it is often impractical.

In this work, we present a PR technique based on an analytical derivative of the vectorial diffraction model that converges to a robust and accurate solution in a pixel-wise manner. Our implementation is very fast thanks to the GPU-optimized implementation using Fast Fourier Transforms (FFT). We provide a flexible software, suitable for both freely rotating emitters and fixed dipoles [27]. Furthermore, our method requires no priors on the pupil plane phase pattern (making it especially suitable for adaptive optics) and, intriguingly, also corrects for experimentally-observed effects, e.g. axial-position dependent lateral shift (also known as wobble), removing the need to calibrate and correct data in post-processing [28]. Our approach is designed to retrieve the phase information from a simple calibration measurement: images of a single bright emitter over a range of focal positions. The retrieved phase can then be applied to volumetric samples, e.g. 3D imaging of mitochondria and microtubule blinking data in live cell imaging.

## 2. Methods

### 2.1 Optical models

An illustration of the optical setup is shown in Fig. 1(a). Briefly, the imaging system consists of a Nikon Eclipse-Ti inverted fluorescence microscope with a Plan Apo 100X/1.45 NA Nikon objective. The emission path was extended with two achromatic doublet lenses (focal length 15 cm) forming a 4-f system. A Liquid Crystal Spatial Light Modulator (LC-SLM) is placed in the conjugate back focal plane (Meadowlark 1920X1152 Liquid Crystal on Silicon) with an EMCCD camera (iXon 897 Life, Andor) at the image plane. The calibration sample consists of a water-covered glass coverslip (Fisher Scientific) with 40 nm fluorescent beads adhered to the surface (FluoSpheres (580/605), ThermoFisher), which is illuminated with 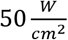, 560 nm light (iChrome MLE, Toptica).

**Fig. 1.**
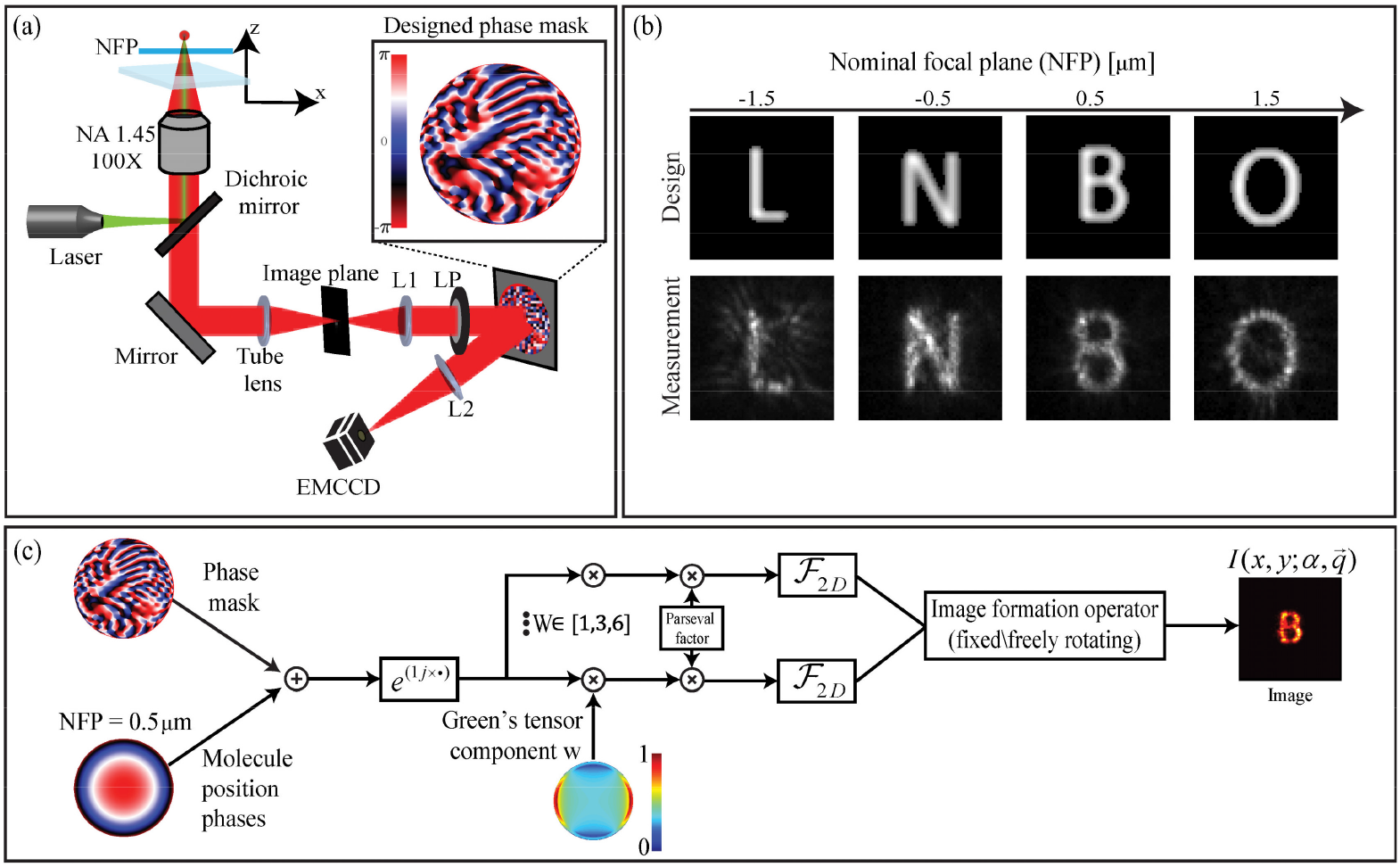
(a) Standard PSF engineering optical setup. The emission from a fluorescent molecule is collected by a high NA oil-immersed objective. A 4-f Fourier processing setup extends the optical path and enables the introduction of a phase mask. (b) (Experimentally measured) Designed phase mask (using our method) which encodes four letter PSFs at different focal plane positions - demonstrates the generality of PSF engineering. (c) Schematic of our simulation model described in section 2.1 where the position of the molecule and the phase mask define the PSF.

Modeling the 3D PSF of the microscope can be done in various methods. A frequently used model is known as the scalar approximation. This model presupposes that the emission pattern of each point source is spherical. This is a good approximation when the objective NA is low, the emitter is rotationally mobile [29], and particularly when imaging emitters far from optical interfaces (beyond near-field effects). Another crucial difference between the vectorial and scalar models is that volumetric imaging far from the coverslip (beyond ~1 wavelength), has an effective NA that is limited by the sample’s refractive index rather than the objective NA. The main advantage of the scalar model is the reduced computational complexity in numerical simulations. Our method implements the more complex vectorial model (in addition to the scalar model) to account for the mentioned effects.

In this section, we discuss freely-rotating emitters for both the scalar and vectorial models. Here, the intensity pattern *I*(*x*, *y*; *α*, *q*) of a quasi-monochromatic source at the image plane is given by [30–32]:

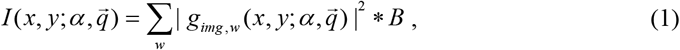

where *α* encompasses the optical parameters required to model the imaging system, including the phase mask, refractive indices, objective parameters, and more. 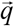 describes the coordinates relative to the coverslip: 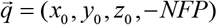; (*x*_0_*y*_0_, *z*_0_) are the emitter’s position coordinates and Nominal Focal Plane (*NFP*) is the objective displacement from focusing on the coverslip. *w* are the electrical field components in the model (Appendix A): *w* = 1 for the scalar model, *w* = 1,2…6 for the full vectorial description and *w* = 1,2,3 or *w* = 4,5,6 for the vectorial model with a linear polarizer, relevant for polarization dependent LC-SLMs. *B* denotes the optical blurring (estimated by a Gaussian kernel with a spatial standard deviation of *σ_B_*). In some models [30,33], an additional filter is added which describes the 3D distribution of emitters for the case of large fluorescent particles. For beads smaller than the wavelength, we approximate these filters by a Gaussian blur to reduce computational complexity.

The electrical field components *g_img, w_* (*x*, *y*) in Eq. (1) are generated by:

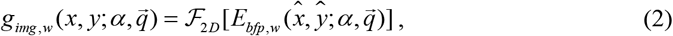

where 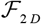 denotes the 2D Fourier transform operator and *E_bfp,w_* represents the electrical field components at the back focal plane given by:

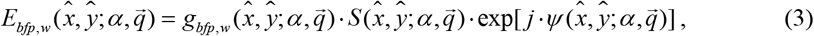

*g_bfp,w_* and *S* are the Green’s tensor components and back focal plane support (aperture), respectively (both are defined in Appendix A). *ψ* is the phase term defined by:

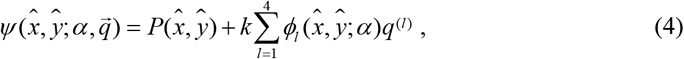

where 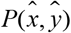 is the phase mask that we retrieve and *q*^(*l*)^ are the elements of 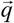. We assume that the phase mask is a single matrix for all the electric field components, i.e. there are no birefringent effects in the back focal plane and we neglect any axial aberrations (which cannot be described by a single phase mask). *k* is the free-space wave number, and 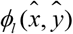 are known phase terms that are associated with the 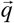 coordinates (Appendix A).

In the next sections, we assume that the optical system is fixed and that we measure a pixelated version of the intensity 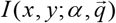. Thus we drop the dependence on the optical system parameters *α*, and keep only the phase mask *P* and the blur kernel std *σ_B_* as free parameters, e.g. 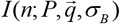, where *n* = 1,2…*N* is the pixel number.

### 2.2 Cost functions

The next step in optimization is introducing the observed z-stack. To do so, a cost function which penalizes the difference between the simulation and measurements is required. Such cost functions can serve in both the PR step and subsequent localization steps. All of the considered cost functions are of the form 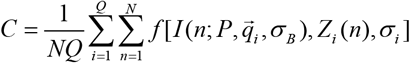 where *Z_i_*(*n*) is the observed image associated with positions 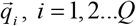, e.g. in the typical case of PR from a z-stack of a bead on the coverslip surface, these correspond to different values of *NFP*.

Common examples of cost functions include the L1 [34] and L2 [24] norms. Other cost functions employ statistical estimation, e.g. are derived assuming Poisson distribution and thus define a negative log-likelihood cost [11,25,35,36]:

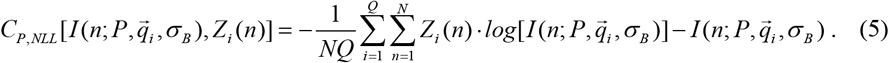

We assume that the intensity distribution 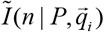 is governed by Poisson statistics with a large mean value, thus a Gaussian approximation is valid (by the law of large numbers) in good calibration samples. In many cases, an additive background term is introduced to the model PSF to better match experimental data [25]. This background noise *bg*(*n*) can be approximated to be white noise, independent from the emitter, with variance of *σ_i_*^2^:

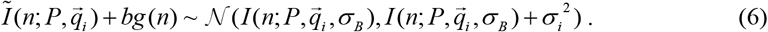

This distribution defines a negative log likelihood cost function:

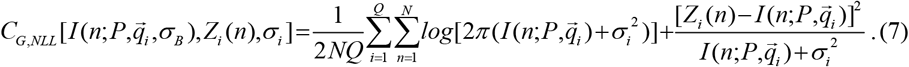

As can be seen in Eq. (7), the resulting cost function is a weighted L2 norm added to a logarithmic term. Unlike *C_P,NLL_*, the cost *C_G,NLL_* comes from a continuous distribution, which is better for derivative calculations. A comparison between the two latter costs is given in Appendix B in terms of their applicability and performance in localization.

### 2.3 Analytical gradient for Phase Retrieval

Phase retrieval can be viewed as a nonconvex optimization problem [37]. In general, such optimization problems pose a challenge as they can have spurious local minimizers, saddle points, and significant sensitivity to good initialization; However, it has been shown that PR can achieve convergence using Stochastic Gradient Descent (SGD) [18].

To facilitate a fast gradient computation, we analytically derive the gradient of the cost function with respect to the pixelated phase mask *P*(*m*), *m* = 1,2…,*M*. In this section we describe the general derivation for dipoles with full rotational mobility (the fixed dipole case is derived in Appendix E). For simplicity, we derive the gradient for a single position 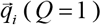. The first derivative in the chain rule (for the form of cost functions defined in section 2.2) gives:

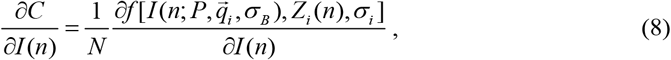

which can be applied to any of the cost functions in section 2.2. The second step is the derivative of Eq. (1) which relies on Wirtinger coordinates [18,38]:

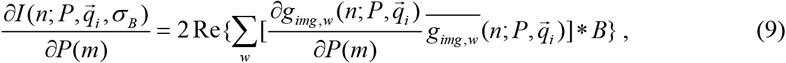

where 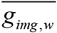 denotes the complex conjugate of *g_img,w_*.

The matrix element 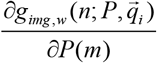 is derived according to Eqs. (2–4):

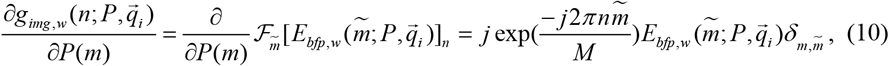

where 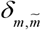 denotes Kronecker’s delta and 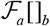 is the Fourier transform over index *a* measured at pixel *b*. The final step is the multiplication of Eq. (8) and Eq. (10):

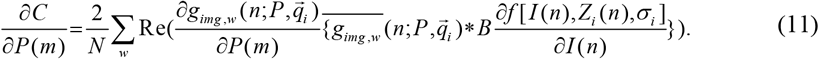

By zero padding Fourier space such that *M* = *N*, the matrix multiplication in Eq. (11) gives a Fourier transform over n (using Eq. (10)):

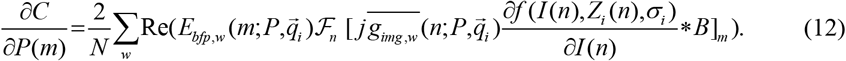

Using Eq. (12), we calculate a single iteration using a few Fourier transforms: the full vectorial model (*w* = 1,2,…,6) requires 12 FFT calculations of size *M* per z position: six to generate the image and six to generate the gradient; critically, a pixel-wise numerical derivative would require 6(*M* + 1) FFT calculations of size *M* per z position (six FFT calculation for a single direction derivative at any pixel and another six to generate the initial image).

### 2.4 Phase mask optimization workflow

Phase retrieval can be used either for *recovering* the pupil plane from measured PSFs, or for *designing* a new phase mask which will produce desired PSFs. Both of these applications amount to very similar procedures. In this section we describe the optimization flow which generates an estimated phase mask from the measured 3D PSF using the previously derived gradient. For mask *design*, the difference is that no denoising or experimental blur correction are required. The first step in the optimization procedure requires the input of a z-stack *Z_i_*(*n*) of length *Q* with associated coordinates 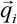. Note that the PR result is the estimation of the phase in the back focal plane, assuming known coordinates 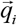. Any mismatch between experiment and assumed coordinates will affect the result. For example, an inaccurate axial position input is compensated by an added defocus correction in the phase mask. Optimally, as many images would be used as possible within the axial range of interest, considering practical limitations, e.g. stage resolution and photobleaching. Next, the background is estimated from the corners of the images to subtract the mean and estimate the noise standard deviation 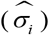.

We initialize the phase mask with an analytical gradient similar to [18]. This improves the convergence rate, because at this point, the simulated images (standard PSF) and measured images are not necessarily correlated. Subsequent *K*_1_ iterations calculate the gradient for a subset of images *via* Adam [39] with a step size of *η* = 3 · 10^−2^. The estimated blur kernel 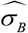 and the noise 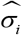 are drawn from a normal distribution in each iteration to reduce the dependence of the solution on the exact estimates. The full algorithm is described in Algorithm 1.

Two optional steps have been included in the software which may improve results in some cases at the cost of additional computation time. The first is fine-tuning the blur kernel (default 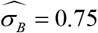) after ⌊1/*K*_1_⌋ iterations. The blur kernel estimation is done using the MATLAB routine *fmincon*. The second option is performing *K*_2_ iterations with a variant of directed sampling [40,41] where the side information is the correlation of the current model with the measured image stack. This can improve results when the used phase mask converges poorly at some focal positions; in this work, we found this step necessary only for the misaligned Tetrapod (Fig. 4. and Appendix D). Future work can include optimization of this final step based on recent advances in adaptive sampling [42], which could further improve convergence.

Importantly, this method is very fast, thanks to the FFT implementation (Eq. (12)): Table 1. Shows the average time required to calculate a single gradient as a function of number of pixels (binning) in the phase mask. The algorithm is orders of magnitudes faster than numerical pixel-wise gradient calculation [24], and much faster than fitting Zernike polynomials, which can take ~30 minutes when implemented using the MATLAB function *fminunc* [34]. The results were timed on a laptop with INTEL i7-8750H CPU and NVIDIA GeForce RTX 2070 GPU. Typical pupil plane recovery (vectorial model with a linear polarizer) from a fluorescent bead z-stack requires 300 iterations over 44000 pixels, using a batch size of 3 z positions, and takes 31.5 seconds on a GPU (Table 1). An equivalent numeric pixel-wise gradient based calculation takes ~9 days (~25,000 times slower).

**Table 1.** Average run time for a gradient calculation using the analytical gradient vs. a numerical gradient.

| Phase mask parameters | Run time GPU analytical gradient | Run time CPU analytical gradient | Run time GPU numerical gradient | Run time CPU numerical gradient |
| --- | --- | --- | --- | --- |
| 175,000 | 0.046[s] | 0.3[s] | 5250[s] | $\sim 35000$[s] |
| 44,000 | 0.035[s] | 0.13[s] | 880[s] | 2640[s] |
| 10,000 | 0.032[s] | 0.032[s] | 200[s] | 195[s] |

#### Algorithm 1: Analytical phase retrieval

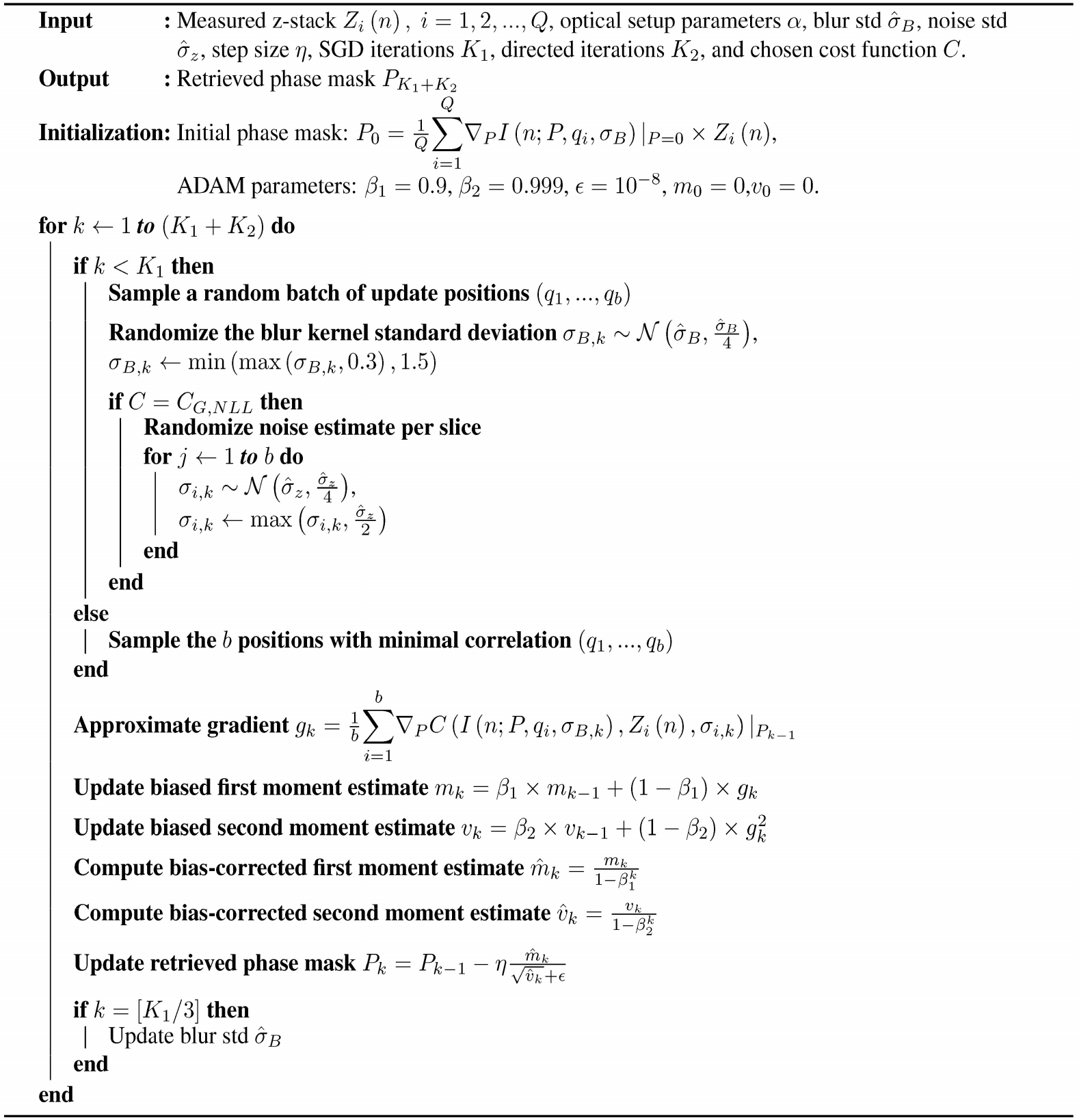

## 3. Results and Discussion

### 3.1 Comparison with existing methods

For benchmarking, we compare our vectorial PR (VIPR) to two alternatives: (1) analytically optimizing (with our method) the scalar diffraction model and (2) adding Zernike polynomials to the initially designed phase mask using the vectorial model [21,34]. Note that using Zernike polynomials requires an additional step to address observed wobble, by fitting the z-stack with the initial guessed mask to center it; this only works when the initial guess is close to the actual mask. Using the described methods, we retrieved phase masks from a z-stacks using a 4 *μm* Tetrapod phase mask [11,43], with one image per NFP (estimated image sum (Signal) of 3.5 · 10^5^[*ADU*] and background 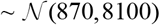). To quantify the performance, we analyzed z-stacks with 10 images per focal position. Robustness of our approach to image noise was tested by computationally degrading the images with a Gaussian-distributed noise (std equal to a multiplier scalar times the estimated std of the noise (Fig. 2(b-d)).

**Fig 2.**
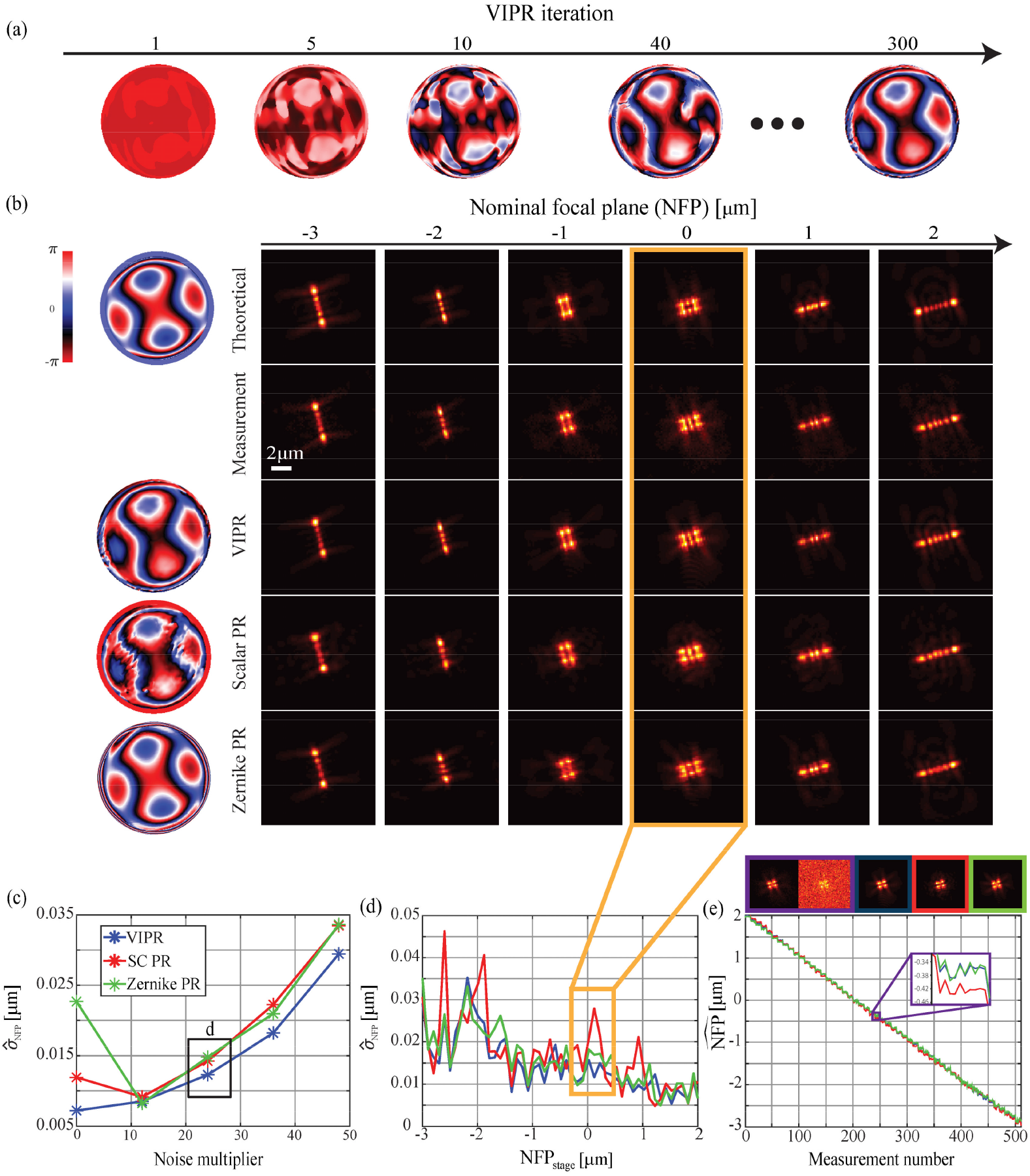
(a) Optimization evolution of the PR mask. (b) First row is the theoretical simulation of the designed phase mask (left) and PSFs (right). Second row shows the acquired measurements when imaging 40 nm fluorescent beads. The next 3 rows show the simulated model using the reconstructed phase masks (left) and PSFs from the tested methods, e.g. VIPR (blue), Scalar PR (using our method with a scalar model-red) and adding Zernike aberration to the vectorial model (green), using the prior knowledge of the designed phase mask. (c) Precision error on the full axial range as a function of added noise and (d) Precision error in each z position at a single noise multiplier (24) of (e) a full axial range acquisition using each of the retrieved phase masks. The orange outline shows a single PSF and MLE fits using each of the phase masks.

While our method deteriorates in precision with added noise, the other methods, interestingly, initially improve after the addition of some noise; this is likely due to our method capturing the subtle details of measured intensity patterns and thus avoids bad localizations at specific axial positions (Appendix C). To test the axial localization bias, i.e. the z accuracy, and the wobble behavior we evaluated the performance of the methods on a short z-stack (bounded by the repeatability of the stage in the two measurements, PR accuracy and noise in both z-stacks). We found that our method incorporates the wobble better and has minimal axial bias (see Appendix C).

### 3.2 Generality to phase patterns

The benefit of pixel-wise methods is their broad applicability to model any aberration, e.g. non-smooth ones, which require many Zernike modes to describe. To test the generality of our method, we imaged z-stacks with 4 different phase mask using the LC-SLM. The first mask consists of a randomly generated combination of low amplitude Zernike polynomials to test the applicability of the method for adaptive optics (Fig. 3(a)). The second case shows the machine learned phase mask which was recently demonstrated to optimally measure dense fields of emitters in 3D (Fig. 3 (b)) [44]. The third test (Appendix D) deals with severe misalignments in the Tetrapod phase mask, which has a large area of unmodulated pixels. Here, for comparison with typically-used algorithms, we also show the results of using centered Zernike polynomials for fitting such an extreme case. The results for the Double-Helix phase mask, which comes from optimization of Gauss-Laguerre basis functions and is commonly used in PSF engineering [45] are also shown in Appendix D.

**Fig. 3.**
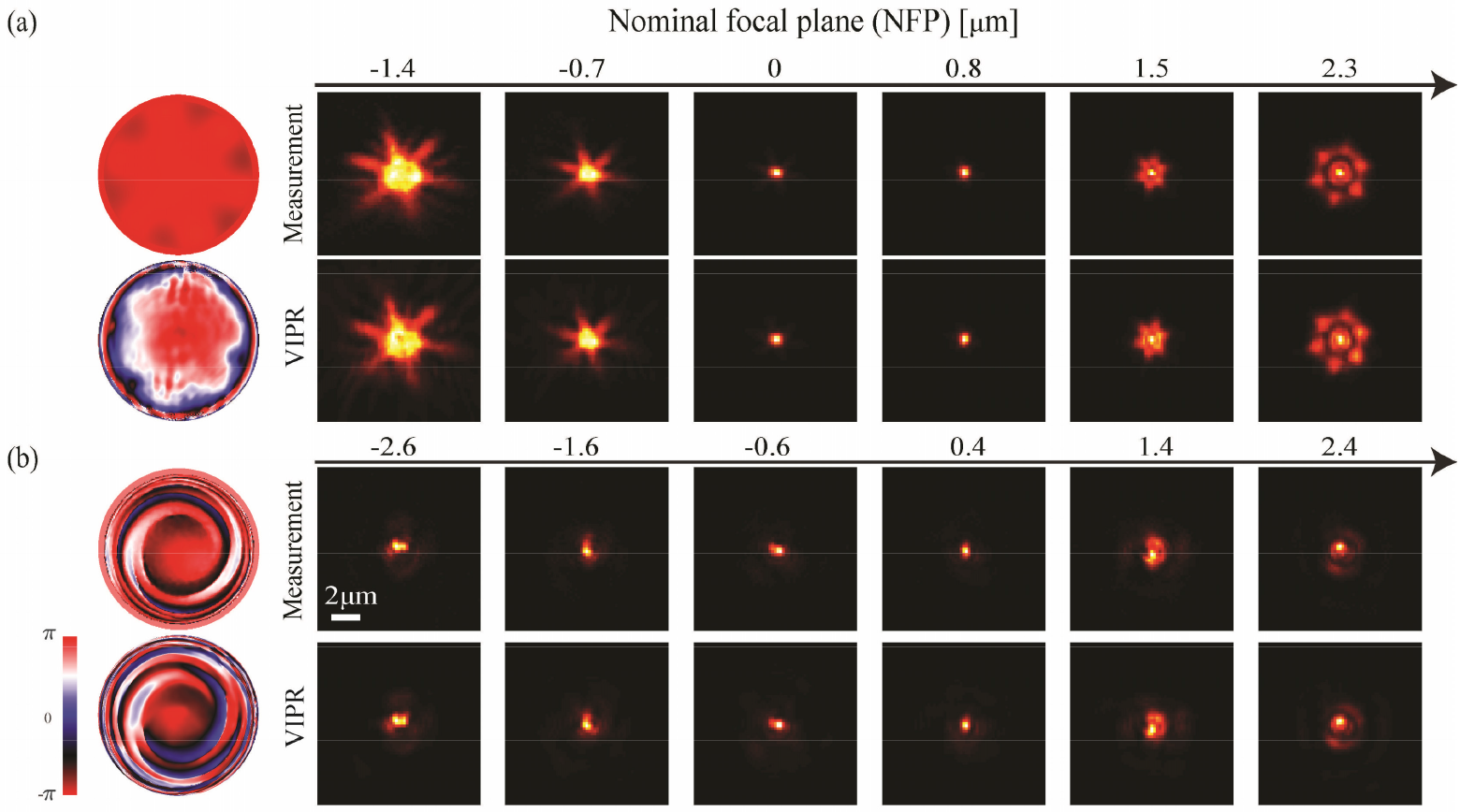
(a) Top row – theoretical phase mask, created from low amplitude combination of random Zernike polynomials and acquired images used to reconstruct the second row using VIPR. (b) Top row – theoretical machine learned phase mask and acquired images used to reconstruct the second row using VIPR.

## 4. Experimental demonstrations of STORM data

Direct stochastic optical reconstruction microscopy (dSTORM) is a technique in which a super-resolution image is reconstructed from localization of sparse fluorescent molecules that are switched on and off (blinking) using high power laser [46,47]. Both experiments (section 4 and Appendix F) were performed using a dielectric 4 *μm* Tetrapod phase mask, which exhibits better photon efficiency than the LC-SLM polarization dependent implementation. PR was performed in a similar way to the LC-SLM data (by removing the polarizer). Appendix F shows a reconstruction of fluorescently labeled mitochondria in COS7 cells, using the deep-learning network which trains on simulations only, adapted from [44]. This demonstrates the viability of our *in-vitro* method in predicting *in-vivo* PSFs.

To show the robustness of our method to suboptimal situations, we used a misaligned 4 *μm* dielectric Tetrapod phase mask to image microtubule blinking data. We show in Fig. 4. the applicability of our method in recovering unusual aberrations, which are otherwise challenging to model. The use of image-based interpolation methods can also address these issues; However, such methods need a good volumetric calibration [48] and are challenging to deploy on a GPU as it requires much more parameters to model the PSF [22]. The PR predictions (using the vectorial model) were used to localize 10K frame of cell data (Fig. 4(a)). Using the localization and an axial range of ±25 *nm* around the desired positions, we created mean experimental PSFs (Fig. 4(c)) to show their matching with our model.

**Fig. 4.**
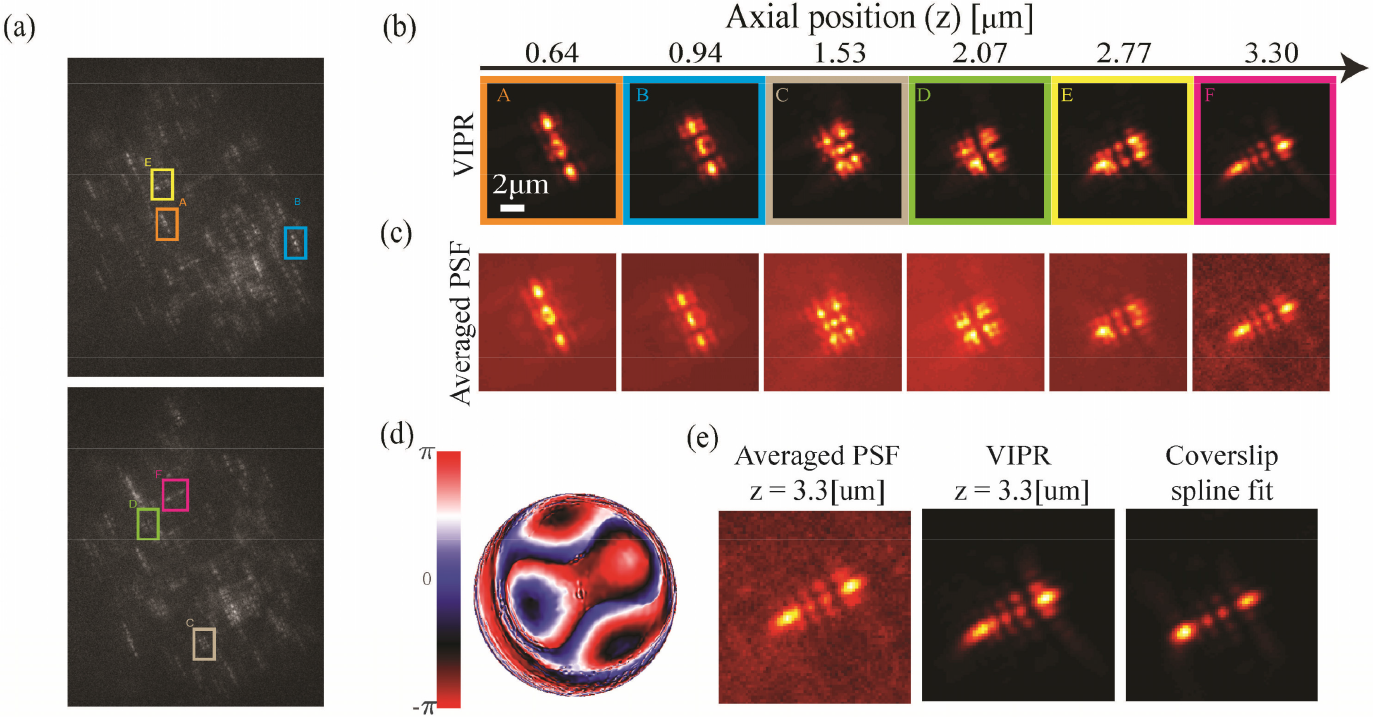
(a) Two example experimental frames taken from a microtubule blinking experiment covering a range of ~3.5 *μm*. Distinct PSFs (A-F) from various axial positions were picked by a localization with deep-learning net that was trained on: (d) the phase retrieved mask (by VIPR) from a coverslip z-stack with FluoSpheres(625/645). (b) Simulated PSFs at the axial position matching to emitters A-F using the PR mask with VIPR. (c) Average PSFs from 10K frames in a range of ±25 *nm* around the stated axial position. (e) Measured PSF in the top of the range with the matching simulation and a spline fit performed on the coverslip data. The difference in shapes shows the importance of incorporating the refractive index mismatch in the optical model as PSFs far from the coverslip experience large deviations from the z-stack data.

## 5. Conclusion

In this work, we have developed and demonstrated an accurate and fast optimization method for PR and PSF design. Our method estimates the optical transfer function in a microscope experiment and provides with an accurate imaging model based on Fourier optics. The optimization uses only basic knowledge of the optical system and a target image set (simulated or measured), with no priors on the phase mask. The derivation is general and can be adopted for many microscopy applications, sample variation (fixed or freely rotating dipoles) and optical processing methodologies (e.g. a polarized LC-SLM, a dielectric phase mask, deformable mirror can be directly modeled with their real pixel size and optical constraints). The performance of VIPR is mainly limited by SNR, the used approximations (e.g. numerical pixelation, aplanatic performance, axially invariant NA, etc.), number of measurements, and accuracy in the knowledge of the optical parameters. The pixel-wise derivation can be adopted to many applications by simply changing the cost function and optimization constraints. Furthermore, this approach can be applied to adaptive optics due to the versatility and speed of our implementation.

## Appendix A

In this section we describe the analytical (far-field) electrical field derivation at the back focal plane which is required in Eq. (3) to calculate - 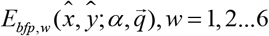. For this section, we assume that there is no phase mask in the Fourier plane (a unit aperture). The source is an emitter located at *r*_0_ = (*x*_0_, *y*_0_, *z*_0_) with a dipole vector 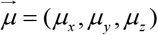. Emitters are immersed in a sample with refractive index *n_m_*, placed above an oil immersion objective with refractive index *n_imm_*. The emission from the source can be described by the far-field Green tensor *G_ff_*(*r*_0_, *r*) [31,32,49], which links between the 3 polarizations of the emitter to the electrical field generated at the far field position 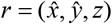:

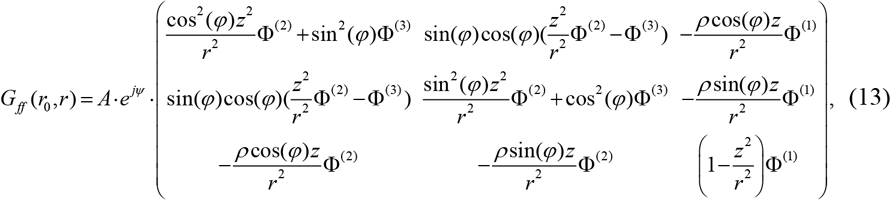

where (*φ*, *ρ*) denote the angular coordinates 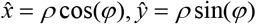. The propagation path is defined by the refractive indices and the propagation angles in the sample medium *θ_m_* and immersion oil *θ_imm_* (connected by Snell’s law). The coefficients Φ^(*v*)^, *v* = 1,2,3 describe the transmission of the different polarizations at the coverslip interface,

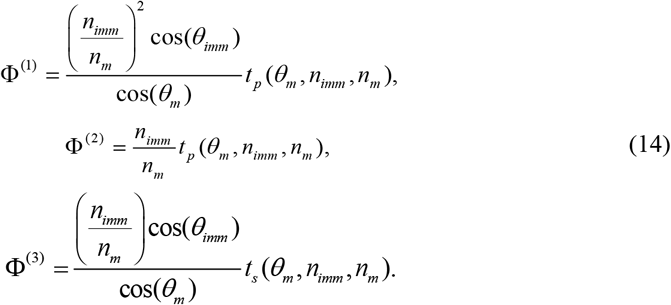

*A* is an amplitude coefficient that can be ignored as the intensity is normalized according to a desired signal. The terms 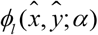 in 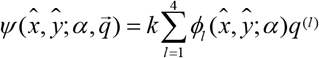 (Eq. (4)) are defined by axial and lateral phases. The lateral phases (*l* = 1,2) are more convenient to describe in the back focal plane using Fourier optics [50] assuming a linear shift invariant system:

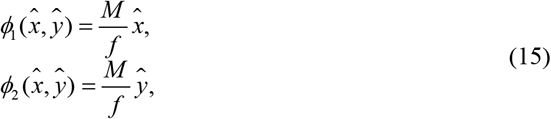

where *M* is the system magnification and *f* is the focal length of the 4-f lenses. The phase induced by the propagation along the optical axis in the different media (*l* = 3,4) under the assumption that this defocus is small compared to the depth of field:

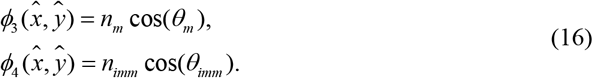

The support of the Fourier plane, 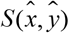 (Eq. (3)), in such systems is usually limited by the objective’s aperture given by Abbe’s sine rule. However, a subtle yet important phenomena can be observed here, usually referred to as Super-Critical Angle Fluorescence (SAF). When the objective’s NA is higher than the refractive index of the sample media, a range of *θ_imm_* propagation angles is available between the critical angle and the NA where cos(*θ_m_*) becomes imaginary. In PR techniques such as this, the molecules are bound to the coverslip and thus the SAF light is measurable.

The last component needed to describe the electrical field at the back focal plane, is the ray rotation matrix which is induced by the objective, assuming a perfect objective lens (full transmission):

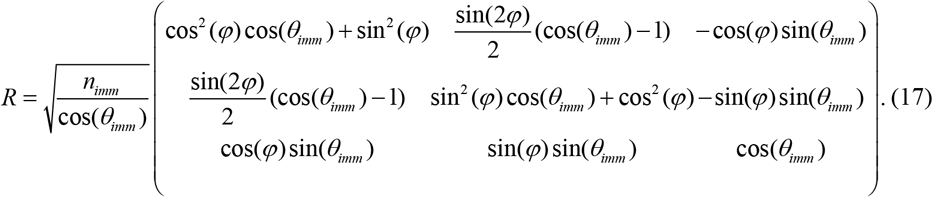

Finally, the electrical field in the back focal plane as we define in Eq. (3) are given by:

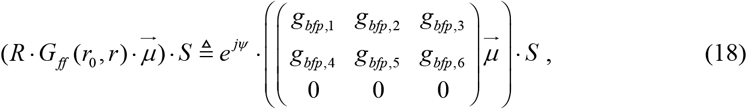

Eq. (18) describes the case of fixed dipoles. Simulating the freely rotating case is simply achieved by super-position (Eq. (2)).

In the case of the scalar approximation, there is only a single component of the electrical field and all the polarization effects are ignored. To relate between the 2 models, the scalar model can be described by 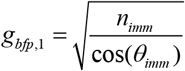:

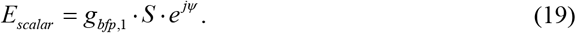

## Appendix B

To compare between the 2 MLE cost functions (Poisson and Gaussian) we used the retrieved Tetrapod phase mask (Fig. 2(a)) to localize the image stack. (Fig. 5(c)). Evaluation of the robustness to noise was achieved by adding artificial noise (similar to section 2.2). We found that the Poisson MLE deteriorates faster than the Gaussian MLE under the addition of artificial noise (Fig. 5(a)) and even under photo-bleaching, which can be seen in Fig. 5(b) where the Poisson MLE has a trend in the precision over the course of a single measurement.

**Fig. 5.**
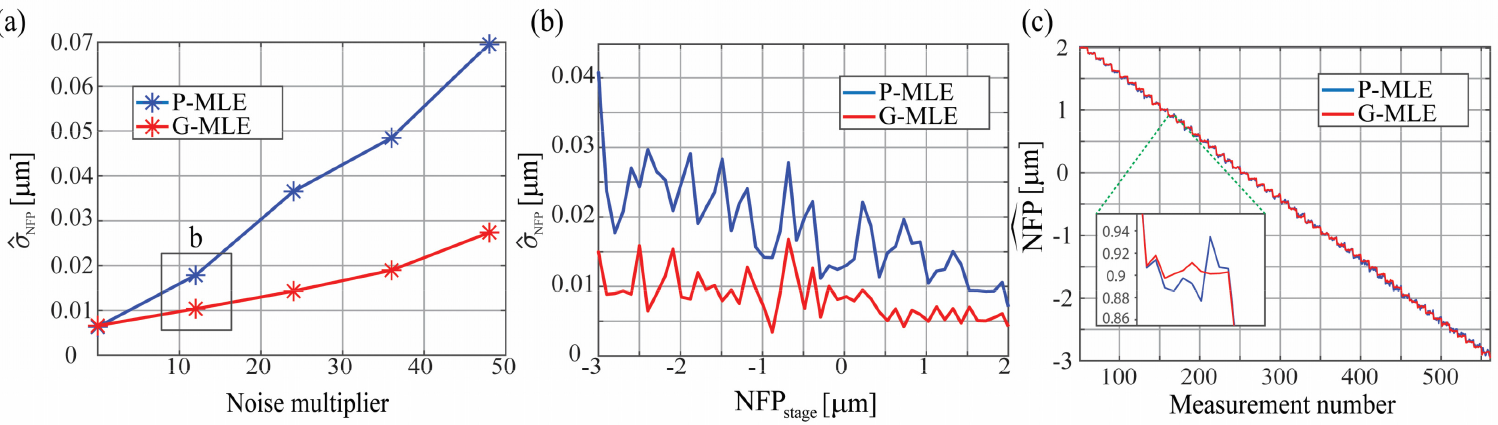
(a) Precision error on the full axial range as a function of added noise and (b) Precision error in each z position at a single noise multiplier of (c) a full axial range localization using the different cost functions.

## Appendix C

This appendix provides with supplementary information to sections 2.3, 2.4 and Fig 2. Figure 6. describes the results of localizing a short z-stack (a second z-stack taken on the calibration bead) with the 3 retrieved phase mask from the described methods in section 2.4. A short z-stack will endure less optical drift and can be used to estimate the accuracy of the PR method. The results are bound by the precision of the stage and the noise in the two experiments (the z-stack which was used for PR and the reconstruction one). Fig. 6(a) shows the difference in the axial position estimation (with MLE fitting) and the stage read where the sole difference is the phase mask. Fig. 6(b-c) shows that the wobble was incorporated into the phase mask using our method (with the vectorial and scalar models) compared to the Zernike PR (green) which shows wobble.

**Fig. 6.**
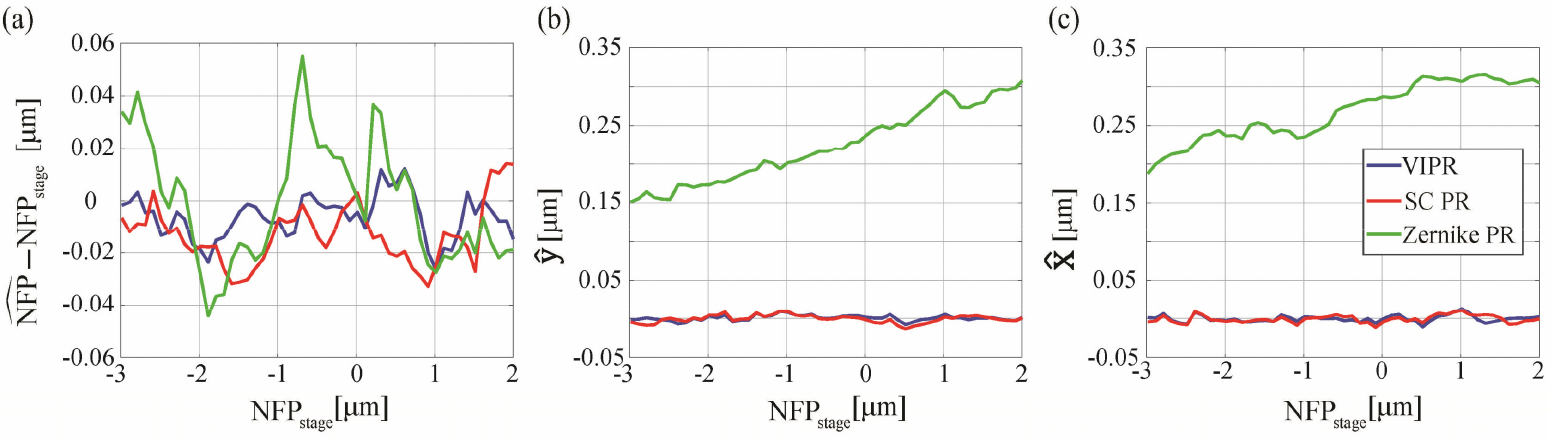
(a) Axial localization mismatch between MLE fit with the three PR masks and the stage reading. (b-c) Lateral localization estimation shows the incorporation of wobble into the phase mask using the pixel-wise approach.

Figure 7. describes the localization results of Fig 2. (b) (without additional noise). From these results, we observe that even a small mismatch in the PSF can lead to large deviations in localization precision at certain positions. These are the positions in which the SAF light produces a sharp point (as it is unmodulated by the phase mask). VIPR incorporates SAF into the derivation and thus suffers less precision penalty in this region.

**Fig. 7.**
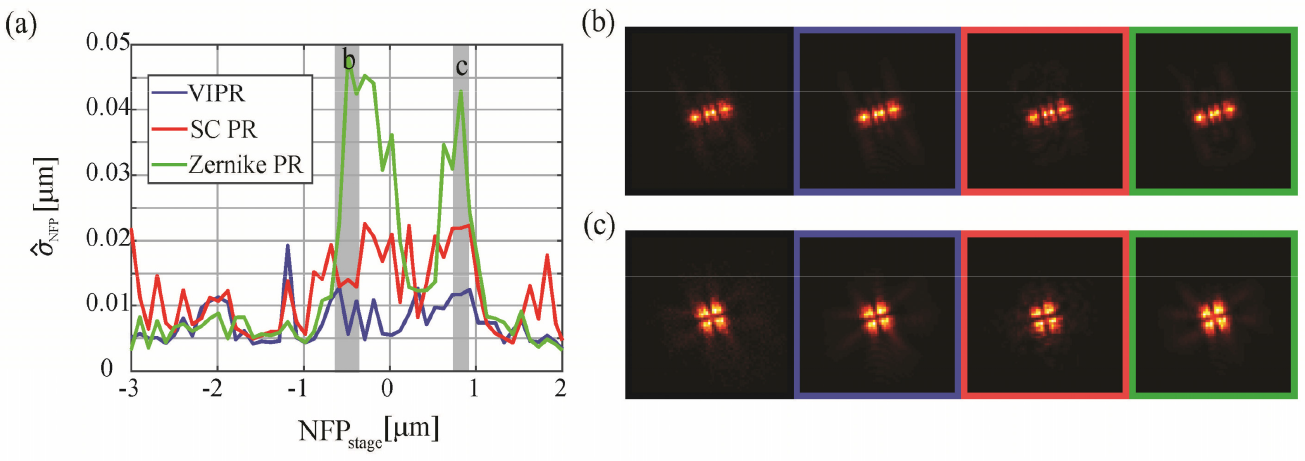
(a) Precision per focal position of the long z-stack with no additional noise, and (b-c) focal positions where the Zernike based method had a large precision scatter due to small differences in the PSF compared to the measurement.

**Fig. 8.**
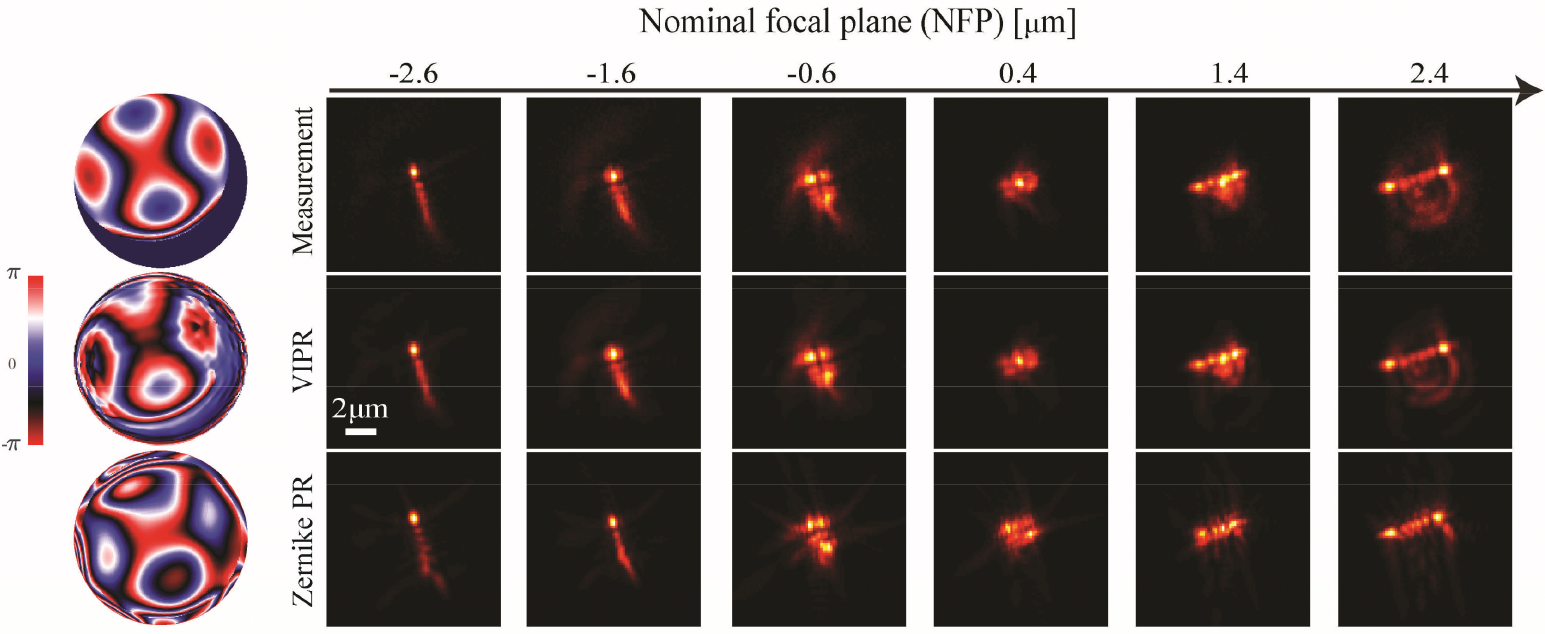
(top row) – Theoretical misaligned Tetrapod phase mask and experimental images. (middle row) - The retrieved phase mask using VIPR and the resulting PSF simulations. (bottom row) - The retrieved phase mask using centered Zernike polynomials and the resulting PSF simulations.

## Appendix D

In this appendix we demonstrate further generality results, in continuation to section 3.2. Fig. 9 shows the retrieval results on a 4 *μm* Tetrapod phase mask which is severely misaligned as described in section 3.2.

**Fig. 9.**
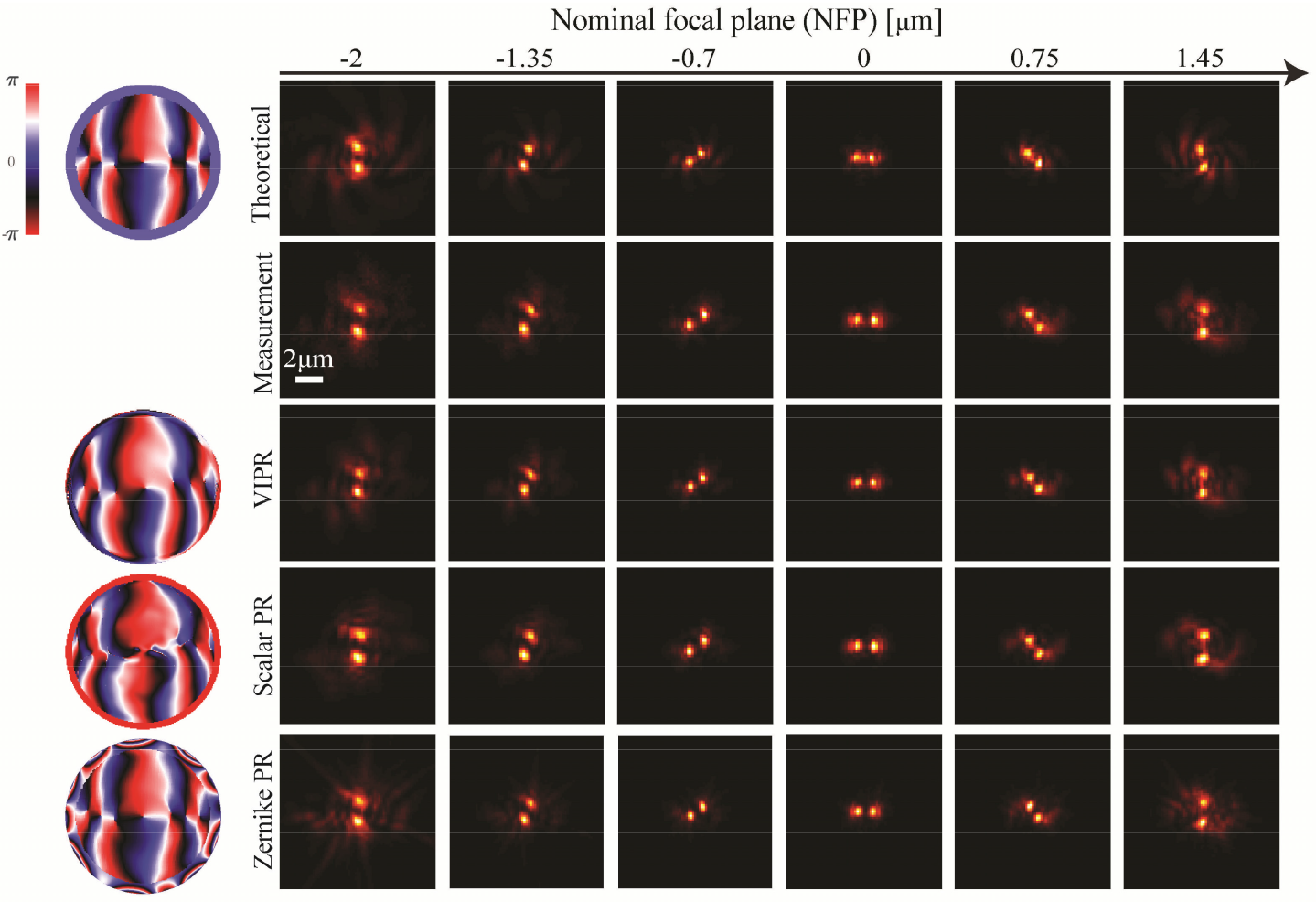
(a) First row is the theoretical simulation the DH phase mask (left) and PSFs (right). Second row shows the acquired measurements when imaging on 40 nm FluoSpheres (580/605). The next 3 rows show the simulated model using the reconstructed phase masks (left) and PSFs from the tested methods, e.g. Vectorial PR (our method), Scalar PR (using our method with a scalar model) and adding Zernike aberration to the vectorial model, using the prior knowledge of the designed phase mask.

Figure 9 shows the retrieval results on a DH phase mask, similar to Fig 2. (a). We note that while the DH is not based on Zernike polynomials like the Tetrapod mask, it can still be corrected using Zernike polynomials if we have the prior knowledge of the theoretical phase mask.

## Appendix E

In this appendix we derive an analytical gradient for the case of a fixed dipole, represented by the vector 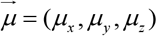. In this case, Eq. (2) is replaced by:

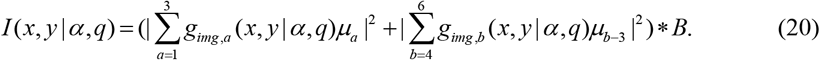

To generate a gradient 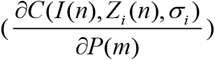 the only change required is deriving Eq. (20) instead of Eq. (2). Thus, Eq. (9) is replaced by Eq. (21):

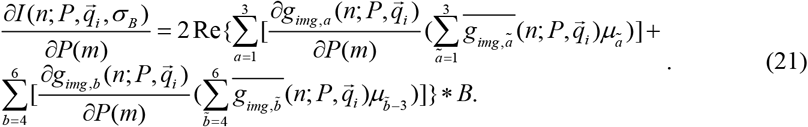

All the subsequent derivatives and the Fourier transform representations are the same as in the freely rotating case. In Fig. 10, we show the theoretical capabilities of this method. We simulated a z-stack of a fixed dipole 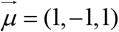 with a linear polarizer and a Tetrapod phase mask. The proposed gradient in this section was used to recover the phase mask.

**Fig. 10.**
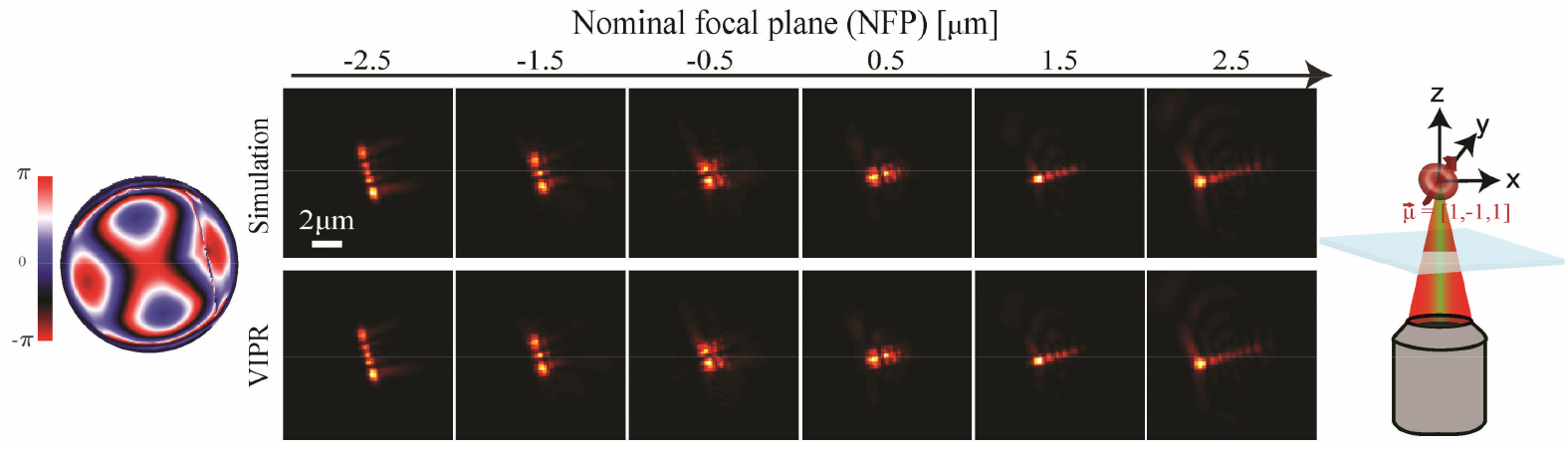
First row is the simulated z-stack of a fixed dipole (right illustration) and a 4 *μm* Tetrapod phase mask, the second row shows the reconstructed PSFs associated with the reconstructed phase mask on the left.

## Appendix F

In this appendix, we describe the sample (section 4) preparation and provide also with a demonstration of VIPR when used on an aligned 4 *μm* Tetrapod phase mask. Sample preparation included cleaning 22X22 mm, 170 *μm* thick coverslips in an ultrasonic bath with 5% Decon90 at 60°C for 30 min. Next, the coverslips were washed with water then incubated in ethanol absolute for 30 min and finally sterilized with 70% filtered ethanol for 30 min. COS7 cells were grown for 24 hours on the coverslips in 6-well plate in phenol red free Dulbecco’s Modified Eagle Medium (DMEM) With 1g/l D-Glucose (Low Glucose) supplemented with 10% Fetal bovine serum, 100 U/ml penicillin 100 ug/ml streptomycin and 2 mM glutamine, at 37°C, and 5% CO2. The cells were fixed with 4% paraformaldehyde and 0.2% glutaraldehyde in PBS, pH 6.2, for 45 min, washed and incubated in 0.3M glycine/PBS solution for 10 minutes. The coverslips were transferred into a clean 6-well plate and incubated in a blocking solution (10% goat serum, 3% BSA, 2.2% glycine, and 0.1% Triton-X in PBS, filtered with 0.45*μm* PVDF filter unit, Millex) for 2 hours at 4°C. The cells were then immunostained overnight at 4°C with either anti TOMM20-AF647 (Fig. 11.) antibody (Abcam, ab209606) or anti-*α* Tubulin-AF647 (section 4) antibody (Abcam, ab190573) diluted 1:230 in the blocking buffer, and washed X5 with PBS.

**Fig. 11.**
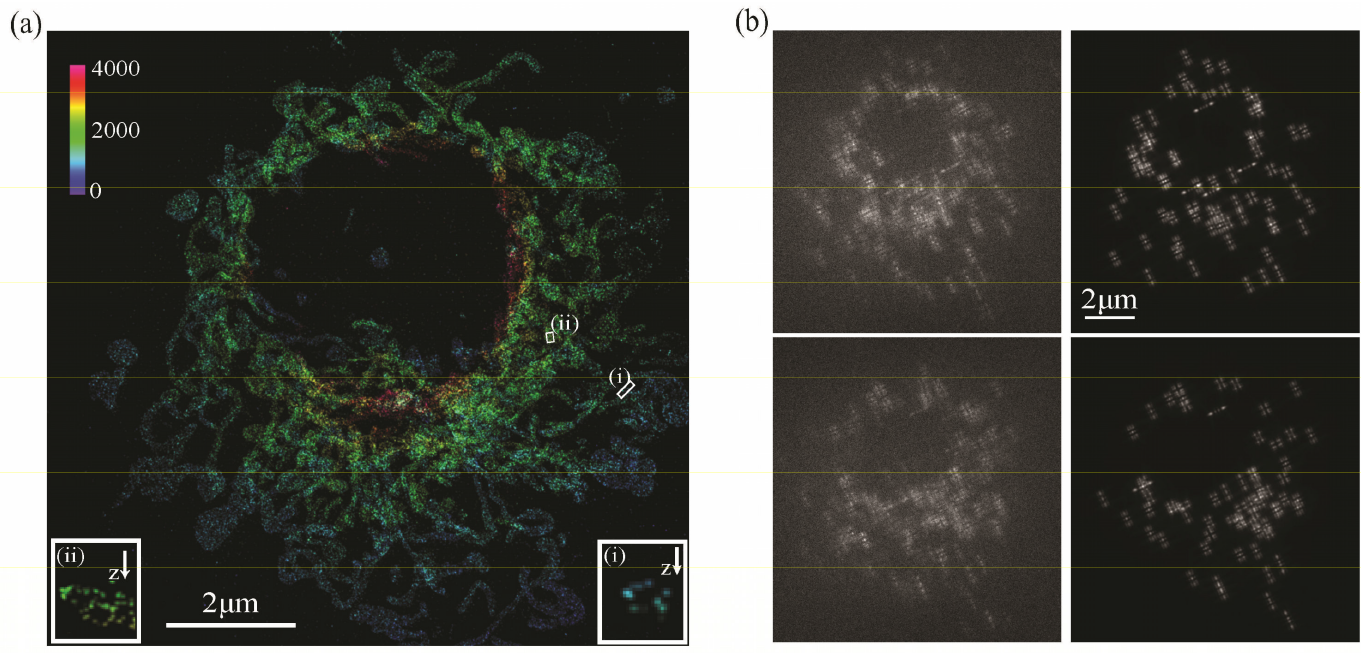
(a) 3D super-resolved image (color encodes depth) of mitochondria in a COS7 cell achieved by training a deep-neural network on simulations of the phase retrieved mask. (i) and (ii) show XZ cross-sections of the white squares, where the hollow shapes of the mitochondria are clearly visible. (b) Two experimental frames (left) and their reconstructions using the phase mask from the PR (right).

For STORM, a PDMS chamber was attached to the glass coverslip containing fixed immunostained COS7 cells and placed in the microscope setup described in Fig. 1(a). The added blinking buffer was freshly made from 100 mM β-mercaptoethylamine hydrochloride, 20% sodium lactate, and 3% OxyFluor (Sigma, SAE0059), modified from [51]. A glass coverslip was placed on top to prevent evaporation. The sample was illuminated by a high-intensity 638nm 2000mW red dot laser module which passed through a 25 *μm* pinhole (Thorlabs). Emission light was filtered through a 500 nm Long pass dichroic and a 650 nm long pass (Chroma). 10K images were recorded on a sCMOS camera (Prime95B, Photometrics) instead of the EMCCD to reduce pixel size and enable a large field of view.

## Acknowledgments

We wish to thank Adam Backer for useful discussions. We thank Navitar Optical Solutions for the donation of a Pixelink camera which was used for system alignment.

## Code availability

The software will be available with the final publication of the paper.

